# Activation microswitches in GPCRs function as rheostats in cell membrane

**DOI:** 10.1101/2020.07.22.216317

**Authors:** Ning Ma, Sangbae Lee, Nagarajan Vaidehi

## Abstract

Although multiple components of the cell membrane modulate the stability and activation of G protein coupled receptors (GPCRs), the activation mechanism comes from detergent studies, since it is challenging to study activation in multi-component lipid bilayer. Using the multi-scale molecular dynamics simulations(50μs), our comparative study between cell membrane and detergents shows that: the changes in inter-residue distances, known as activation microswitches, show an ensemble of states in the extent of activation in cell membrane. We forward a rheostat model of GPCR activation rather than a binary switch model. Phosphatidylinositol bisphosphate (PIP2) and calcium ions, through a tug of war, maintain a balance between the GPCR stability and activity in cell membrane. Due to the lack of receptor stiffening effects by PIP2, detergents promote more transitions among conformational states than cell membrane. These findings connect the chemistry of cell membrane lipids to receptor activity useful to design detergents mimicking cell membrane.

## Introduction

G protein coupled receptors (GPCRs) are seven helical transmembrane (TM) proteins that transduce a variety of cell signaling pathways. Agonist binding leads to coupling of the receptor to trimeric G proteins which initiates a cascade of signaling events resulting in intracellular biological response. Pioneering structural studies on agonists and G proteins bound GPCRs using crystallography, electron microscopy, NMR and DEER techniques have shown that GPCRs are inherently dynamic proteins adopting multiple conformations, such as inactive, active intermediate and G protein coupled fully active conformational states (1–8). These studies have highlighted the structural changes in GPCRs associated with the different conformational states leading to the activation of GPCRs. The dominant structural changes involve movement of transmembrane helices 5, 6 and 7 (TM5, TM6 and TM7) as well as changes in the intracellular loops. Comparison of three-dimensional structures of active state to that of inactive state of several class A GPCRs have shed light on the inter-residue distances that show distinct changes upon activation. These inter-residue distances are called “activation microswitches”(9–11). The activation microswitches are thought of as binary “on or off “ switches, a concept that stems from analysis of static structures. All these structural studies were conducted in detergent micellar solution or in nanodiscs containing 1-palmitoyl-2-oleoyl-sn-glycero-3-phosphocholine (POPC) lipids (12–15). On the other hand, multiple studies have shown that different lipids in the cell membrane play an active role in modulating the activity of GPCRs(15–22). It is also known that cations such as Ca^2+^, and Mg^2+^ play an important role as allosteric modulators of GPCR activity(23–26). NMR studies of β2-adrenergic receptor in detergent micelles compared to reconstituted high density lipoprotein environment showed that the basal activity and exchange rates between inactive and active conformations is higher in detergents and in phospholipid bilayer containing HDL particles compared to cell membranes(15, 27, 28).

Although we know that the cellular environment has a definitive effect on GPCR stability and activity, and even the selective coupling of G proteins by GPCRs (19), there is a serious gap in our understanding, at an atomic level, of the effect of the chemical nature of different lipids and cations on the GPCR conformation dynamics. More importantly, the spatiotemporal component (the persistence time) of lipid-receptor contacts that play an important role in activation mechanism has not been considered while analyzing static structural contacts with lipids. In summary, the similarities and differences of the GPCR activation mechanism in cell membranes compared to detergents is unknown. The effect of cell membrane components on the activation microswitches is also poorly understood. An understanding of these fundamental phenomenon would (i) lead to insights into GPCR dynamics and activation in physiological cell membranes that is critical to cell signaling and (ii) allow us to design detergents that mimic cell membrane behavior more closely, or add lipids that modulate GPCR activity in cells, to biophysical studies conducted in detergents or nanodiscs. Studying GPCR dynamics in cellular environment is challenging experimentally. However, multi-scale Molecular Dynamics (MD) simulations offer a suitable tool to map the conformation ensemble of different conformational states of GPCRs and to probe the role of multiple lipid and cation components of cell membrane on GPCR dynamics.

In this study we have used a combination of coarse grain MD (30 μs) followed by 20 μs of all-atom MD simulations in a hierarchical workflow, to uncover the mechanisms of how different lipids in the cell membrane affect the dynamics and conformation ensemble of GPCRs in different conformational states. For comparisons, we have also performed MD simulations in detergent micelles that is often used to study GPCR activation mechanism. The MD simulations were performed on human adenosine A_2A_R receptor in its agonist bound inactive (R), active intermediate (R’) and miniG protein bound fully active (R*.G) states in cell membrane mimicking bilayer. We chose A_2A_R for this work due to availability of crystal structures in all the three above mentioned conformational states. Our results highlight the fact that not only the nature of lipid interaction with the receptor, but also the *temporal persistence of these interactions* during the dynamics in cell membrane conditions, play a critical role in modulating the receptor flexibility and activity. We show that the detergent environment increases the flexibility of the TM helices and therefore promote transitions between the inactive and active-intermediate state when bound to an agonist. This is in stark contrast to the cell membrane environment especially Phosphatidylinositol bisphosphate (PIP2) and calcium ions that together maintain a balance between the receptor flexibility and stability and therefore limit the transitions between inactive state and active intermediate states.

Analysis of static structures of the inactive and active state of several GPCRs showed that the activation microswitches show a binary on and off switch like behavior (9–11). Our dynamics study that takes the temporal persistence of activation microswitches into account, shows that they behave like *rheostats* in cell membrane (showing different degrees of activation in an ensemble of states) rather than binary on and off switches. The combination of microswitches activated in cell membrane are different than those activated in detergents during the early events of activation. Additionally, the extent to which the microswitches are activated are different in cell membrane and detergents. Taken together, our work provides mechanistic insights into the role of the chemistry of lipids in modulating the GPCR dynamics that can be used for designing detergents that better mimic the cell membrane.

## Results

### Multi-scale MD simulations in mixed lipid bilayer

Previous coarse grain MD simulation studies on A_2A_R receptor in various conformational states showed accumulation of GM3, PIP2 and cholesterol close to the receptor depending on the conformational state of the receptor(20). However, there is very little knowledge on the atomic level mechanism of these interactions and the effect of the chemistry of the lipids and cations, on GPCR conformational dynamics in cell membrane compared to detergents. To study the effect of multiple lipids on the GPCR conformation ensemble, we used multi-component lipid bilayer to mimic the cell membrane. The outer leaflet of the membrane bilayer consists of POPC, DOPC, POPE, DOPE, sphingomyelin(Sph), ganglioside(GM3) and cholesterol in the ratio of 20:20:5:5:15:10:25, while the inner leaflet contains POPC, DOPC, POPE, DOPE, POPS, DOPS, phosphatidylinositol 4,5-bisphosphate(PIP2), and cholesterol in the ratio of 5:5:20:20:8:7:10:25 and all simulations were neutralized by adding 0.15M of NaCl or CaCl_2_ (see Fig. 1A and Table S1). Since this mixed lipid bilayer mimics the cell membrane, we refer to this as “cell membrane” hereafter in the paper. We started the coarse grain MD simulation from the crystal structures of the agonist 5’-N ethylcarboxamidoadenosine (NECA) bound human adenosine receptor A_2A_R in the inactive state R, the active-intermediate state R’ and the mini-G-protein bound fully active state of the receptor R*.G in the cell membrane model (3, 9, 29) (see Methods for more details). We performed 30μs of coarse-grained MD simulations to obtain an equilibrated multi lipid cell membrane mimic system. We extracted the receptor conformations in all the three states from coarse grain simulations and converted them to all-atom models (see Methods) and performed all-atom MD simulations. The list of systems simulated in this study, the notations used to represent different conformational states, and other details of these systems are given in Table S2.

**Figure 1.**
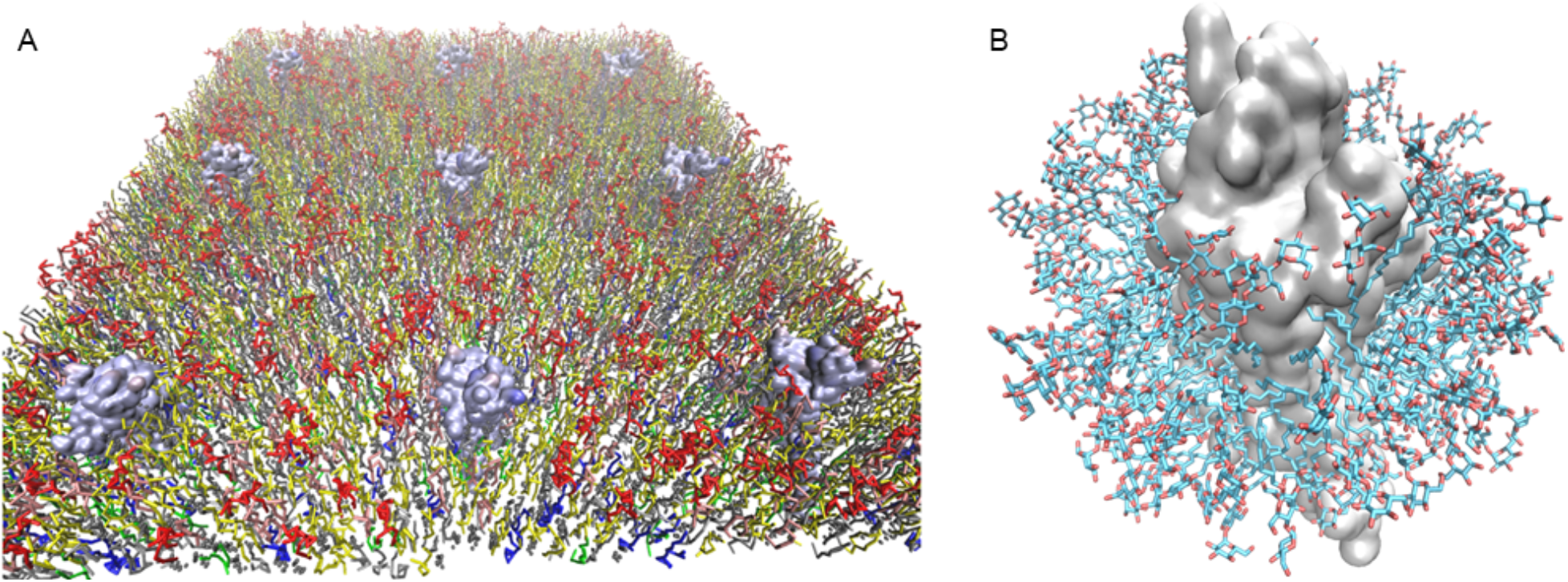
**A.** Starting model of the cell membrane consisting of multiple lipids used in the coarse grain simulations and all-atom molecular dynamics simulations. The colors denote different types of lipids. The surface rendering are of the A_2A_R crystal structures in the agonist NECA bound inactive state R, NECA bound active intermediate state R’ and mini Gs protein and NECA bound fully active R*.G conformation state. **B.** Starting model of the DDM detergent micelle with A_2A_R used for MD simulations.

### MD simulations in Detergents

To compare the effect of the detergent environment on receptor dynamics, to that of cell membrane we also performed all-atom MD simulations on NECA bound R, R’ and R*.G states of A_2A_R in DDM detergent micelle (Fig. 1B) using procedures described in the Methods section. For comparisons, we have also used analysis of our previous MD simulations using branched detergent Lauryl Maltose Neopentylglycol, LMNG(30).

### Agonist binds with higher affinity in cells compared to detergent micelles

Binding affinity of the agonist NECA is an order of magnitude stronger (Kd = 2.4 ± 0.1 nM)(31) in cell based measurements compared to that in DDM detergent solution (23.4 ± 0.11 nM) (32). This translates to a difference of 1.4 kcals/mol in free energy at 310K. Using free energy perturbation alchemy methods (Bennett Acceptance Ratio method – see Methods section) we computed the binding free energy of NECA in R, R’ and R*.G states both in cell membrane and in detergent DDM micelles. The calculated binding free energy of NECA is 2.1 kcal/mol and 0.8kcal/mol better in the cell membrane compared to DDM detergent micelle in the inactive R and active-intermediate R’ states respectively with no difference in the R*.G state (Fig. 2A). We analyzed the structural basis for this enhanced agonist binding affinity for the R state in the cell membrane compared to detergents. We identified the receptor ligand interactions that cause this difference in binding free energies. More importantly, even for the receptor ligand interactions that are common to both detergent and cell membrane environments, the difference in their “persistence frequencies” contribute to the difference in binding energies. The persistence frequency of a residue-residue contact or a receptor-ligand contact is defined as the percentage of MD snapshots that contain that contact. The list of receptor-NECA contacts and their persistence frequencies in the NECA bound R state in cell membrane and in DDM detergent are shown in Fig. 2B (see also Fig. S2). The persistence frequencies of the NECA-receptor contacts are higher in the cell membrane environment compared to the detergent environment for the R state. This shows that the cell membrane components tighten the binding site residue-NECA contacts even in the inactive R state conformation similar to that of the binding site of NECA in the R’ state. This finding demonstrates the importance of dynamics simulations in cell membrane conditions to recapitulate the observed differences between cell membrane and detergents. The agonist NECA shows higher flexibility in the agonist binding site in the R state in detergent environment compared to cell membrane environment as shown by the 3 distinct conformations in detergent shown in Fig. 2C. The receptor also shows higher flexibility in the R state around the agonist binding site, in detergent environment compared to cell membrane environment as seen in the heat map of flexibility shown in Fig. 2C. This shows that the lipids in the cell membrane have a pronounced effect on agonist binding even in the inactive state compared to detergents. Next we identified the lipid components of the cell membrane that bring about this favorable effect on agonist binding affinity in the extracellular ligand binding region in the inactive state.

**Figure 2.**
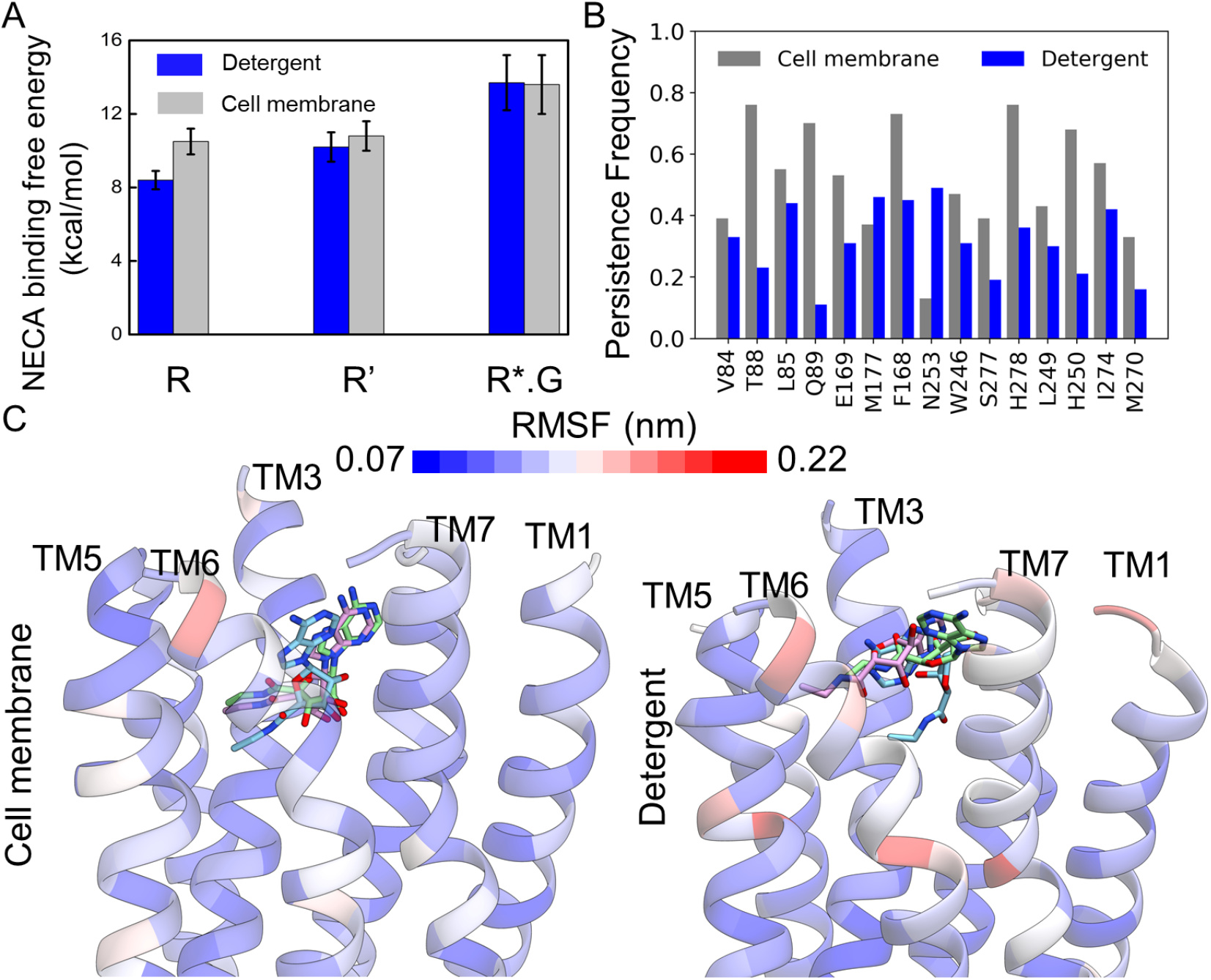
Structural basis for the higher binding affinity of NECA in cell membrane compared to detergent DDM. **A**. Calculated binding free energy of NECA in R, R’ and R*.G conformational states of A_2A_R shown with the respective standard deviations. The free energies were computed using free energy perturbation - BAR method (see Methods section). **B.** Persistence frequency of the NECA-residue contacts in cell membrane (grey) and in DDM detergent (blue). The persistence frequency is the percentage of MD snapshots that contain the NECA-residue contacts. **C.** The conformational flexibility of the agonist NECA in the R state in cell membrane and in detergent. Top 3 representative structures from clustering analysis of NECA are shown. The representative structures of NECA from cell membrane simulations are shown in purple (51%), blue(22%) and green (7%). The respective population of these agonist conformation clusters are given within parenthesis. The representative structures of NECA and the population of the conformation clusters from the detergent DDM simulations are shown in purple (43%), blue (15%), green (11%). The population of the top three occupied clusters of NECA are shown in parenthesis. The RMSF values that measure the flexibility of the Cα atoms of the residues in the NECA binding site, are shown in a heat map overlaid on the A_2A_R inactive R state structure. Blue to red color indicates increasing flexibility of the receptor.

### GM3 tightens the ligand binding site in the extracellular leaflet of the cell membrane

GM3 is present in the outer leaflet of the cell membrane. Its polar head group consists of sugars and has a long hydrophobic tail. During the simulations of all the three conformational states of A_2A_R in cell membrane, we observed that the hydrophobic tail of GM3 binds in exterior crevices between the extracellular regions of TM6 and TM7. The polar head groups interact with the residues G152, K153 and S156 in the extracellular loop2 (ECL2). Additionally, residues in both ECL1 and ECL2 show persistent hydrogen bond interaction with the head group of GM3 as shown in Fig. 3A. Gangliosides have been shown to regulate GPCR function and coarse grain MD simulations have shown that they interact with extracellular loops(33, 34). As a result of these persistent interactions of GM3 with ECL2 residues, we observed a large movement in ECL2 (4 to 8Å) that covers the NECA binding site as shown by the grey arrow in Fig. 3B (and Fig S3B). Such interactions of DDM with ECL2 residues were not observed in the detergent micelles (Fig. 3A) and the ECL2 actually shifted 3Å away from ligand binding pocket (Fig. 3B blue arrow). The residues G152, K153 and S156 in ECL2 showed stable interactions with GM3 head group in R state as shown in Fig. S3B. Calculation of persistence frequency (percentage of MD snapshots that show contact between GM3 and residues in ECL2) of GM3 with these residues show that the GM3 contacts with these residues is most persistent in the R state compared to R’ and R*.G state as shown in Fig. S3C. The closing of ECL2 over the NECA binding site leads to contraction of the volume of the NECA binding site in cell membrane simulations as shown in Fig. 3C. The average volume of ligand binding pocket calculated using the MD snapshots from the most populated conformation cluster decrease from 333.4 Å^3^ in DDM to 279 Å^3^ in cell membrane. The cell membrane components such as GM3 constrain the structural elements around the NECA binding site thus leading to improved interactions with the receptor even in the inactive state. Song et al(20) reported the effect of GM3 on gating the on and off rate of agonist using the persistence of contact of GM3 with the receptor calculated from coarse grain MD simulation study. Here, from the atomistic level simulations we calculated the binding free energy of NECA in cell membrane and in detergents. We also show how GM3 enhances the receptor-agonist interactions in the binding site by tightening these interactions.

**Figure 3:**
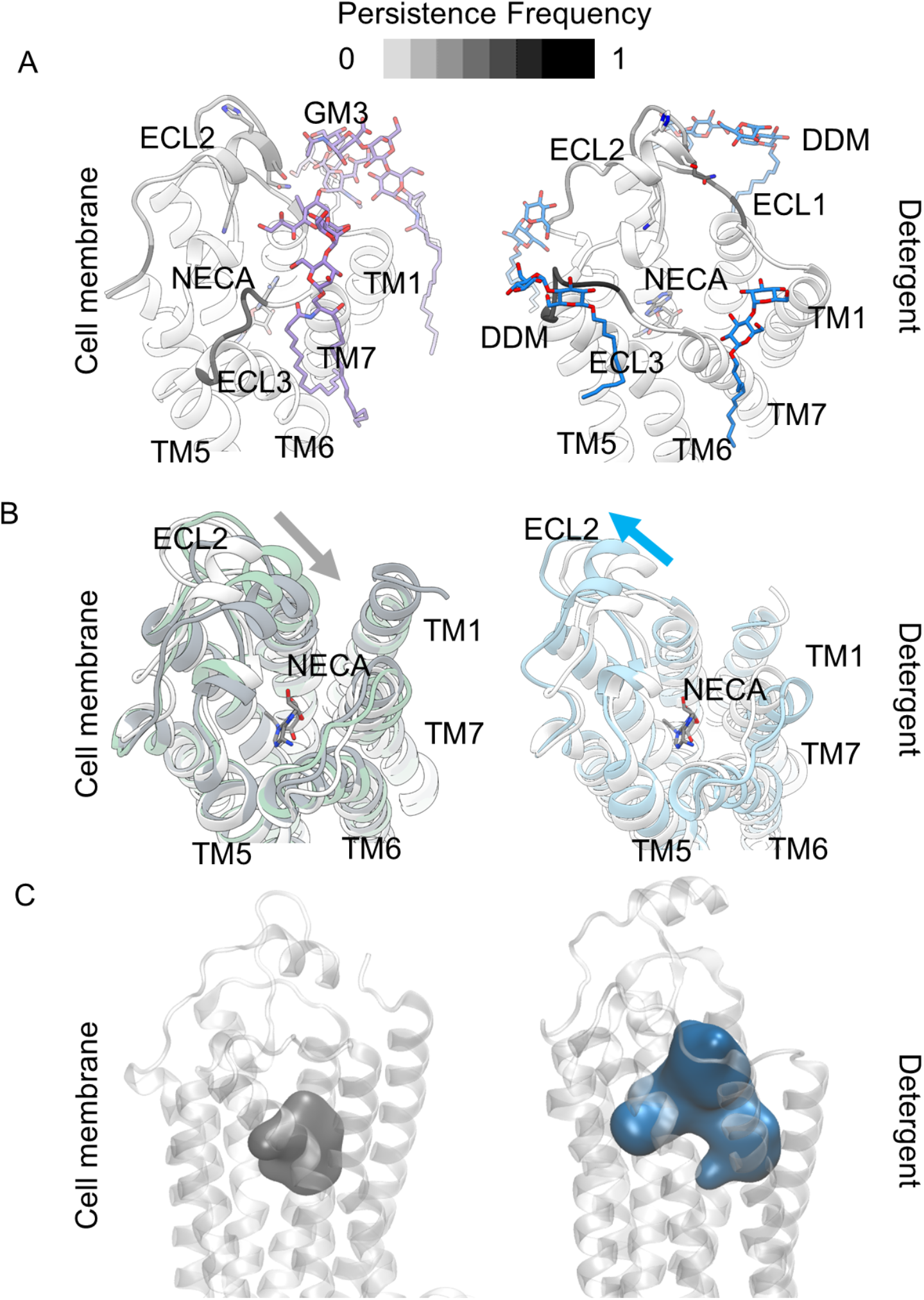
GM3 pulls the ECL2 to cover and shrink the agonist binding site. **A**. The persistence frequency of NECA-residue contacts in the extracellular loops ECL1, ECL2, ECL3 in the R state with GM3 is shown as a heat map on the structure of R state of A_2A_R in cell membrane and in DDM detergent. **B**. Overlay of representative structures extracted from the top two populated conformation clusters extracted from the MD simulations of the inactive R state in cell membrane and in DDM detergent. The starting structure of the simulations is shown in white in these two figures, and the representative structure from the most populated conformation cluster are shown in grey (cell membrane) and cyan (detergent) color. The structure shown in green is the representative structure from the second most occupied cluster in cell membrane. Both these conformations show a closing of ECL2 over the NECA binding site which is not seen in detergent. **C**. The NECA binding pocket shrinks in volume, caused by GM3 closing the ECL2 over the ligand binding site and enhancing the NECA-residue contact frequency for the R state. The ligand binding site volume is larger in the DDM environment.

### Cell membrane environment constrains the conformational flexibility of GPCRs compared to detergents

The movement of transmembrane helix TM6 away from the core of the receptor and movement of TM7 towards TM3 and TM5 has been shown to be major conformational changes in the active state of the receptor compared to the inactive state (35). To assess the extent of conformational sampling and flexibility of the receptor in the intracellular regions, we calculated the C_α_-C_α_ distances between residues R102^3.50^-E228^6.34^ on TM3 and TM6 and between R102^3.50^-Y288^7.53^ on TM3 and TM7 for every snapshot in the MD simulation trajectories. We projected the snapshots from MD simulations in cell membranes and in detergents in these two distance space, as shown in Fig. 4A and 4C for the R, R’ and R*.G states in cell membrane and detergent respectively. In the NECA bound R state, the receptor conformation ensemble shifts away from the starting crystal structure in the R state and the transitions between R and R’ states are more populated in the DDM environment than in cell membrane environment (Fig. 4C). On the other hand, there are no transitions between the R’ and R*.G states observed within the time scale of the all-atom MD simulations in both cell membrane and detergent environment. The spread of the receptor conformation ensemble in cell membrane is more constrained in each R, R’ and R*.G states (Fig. 4A) compared to the detergent DDM environment (Fig. 4C). As detailed in the next section, this is due to the charged lipid PIP2 present in the inner leaflet of the cell membrane accumulating close to the basic residues (Arg/Lys) in the intracellular regions of the receptor in all the three conformational states.

**Figure 4:**
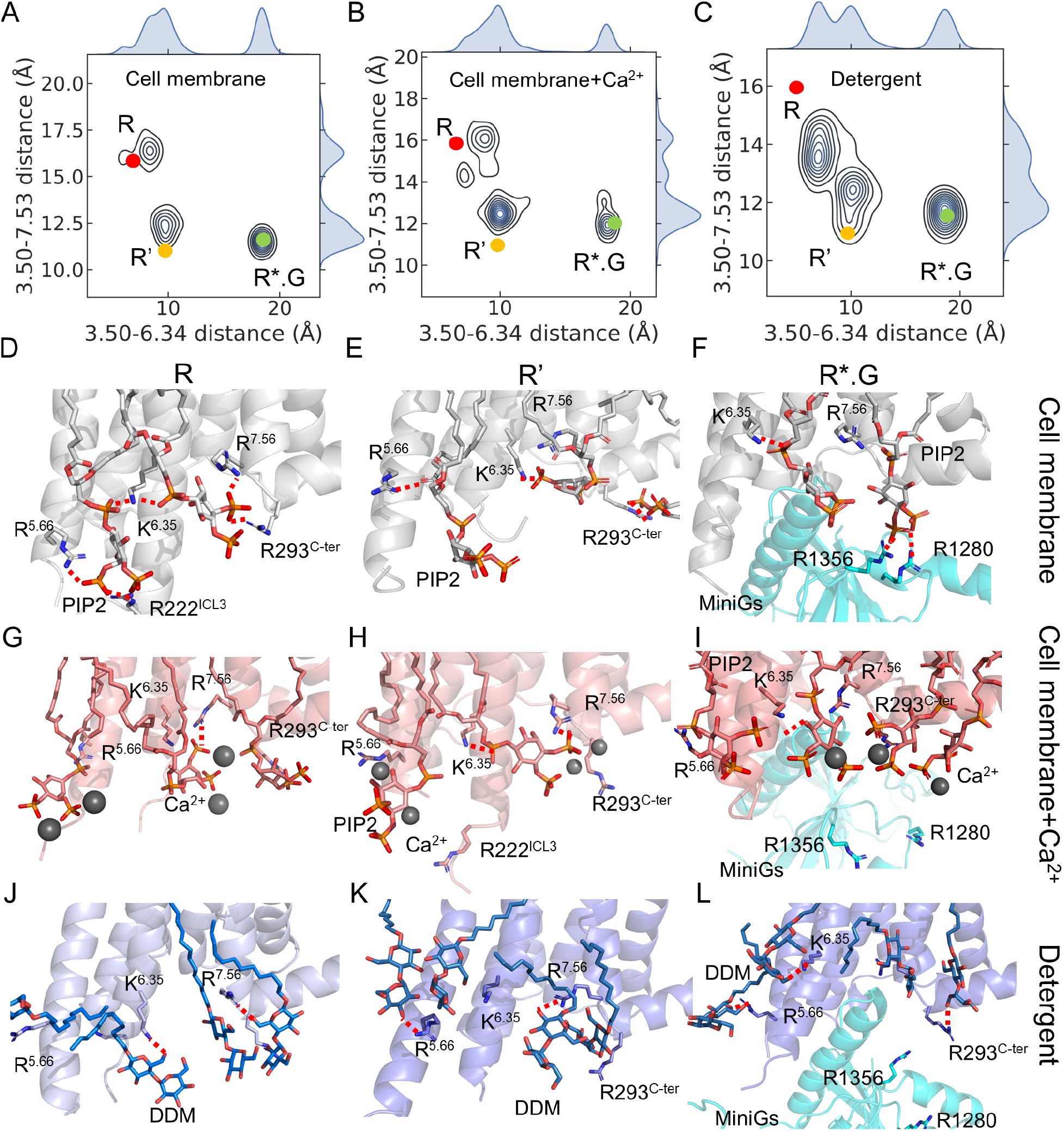
PIP2 and intracellular calcium modulate receptor flexibility and activity. **A-C.** The extent of receptor conformation sampling of NECA bound A_2A_R in R, R’ and R*.G states in three different environments, namely, the cell membrane(A), cell membrane with calcium ions (B) and DDM detergent (C). The MD snapshots are projected on inter-residue distance measures. The distances are measured between the C_α_-C_α_ atoms of the residue pairs R102^3.50^-E228^6.34^ on TM3 and TM6 and between R102^3.50^-Y288^7.53^ on TM3 and TM7. These two distances in the starting crystal structures of A_2A_R in R, R’ and R*.G states are shown as red (R), yellow (R’) and green (R*.G) dots respectively. **D-F**. PIP2 forming bifurcated salt bridge interactions with the Lys/Arg in the intracellular region of A_2A_R, bridging the TM helices, in cell membrane conditions in the inactive R state (D), in the active-intermediate R’ state (E) and in the miniG protein bound fully active R*.G state (F). PIP2 is shown in grey and orange sticks representation. **G-I**. Calcium ions (shown in black spheres) interactions with PIP2 yanks the PIP2 away from the Lys/Arg residues in the receptor R state (G), R’ state (H) and R*.G state (I). Calcium ions coordinate with PIP2 freeing up the TM helices thus promoting transitions between R and R’ state seen in figure B. **J-L.** DDM forms weak hydrogen bonds with Arg/Lys in the intracellular region in the R state (J), R’ state (K) and R*.G state (L). Therefore, the receptor is more flexible in detergents.

### PIP2 reduces movement of TM helices in the intracellular regions

PIP2 accumulates close to the receptor in all the three R, R’ and R*.G conformational states (Fig S1B). In the R and R’ states, as shown in Figs. 4D and 4E, PIP2 forms salt bridge interactions to the Arg/Lys residues in the intracellular region of TM5, TM6 and TM7. These bifurcated salt bridges of PIP2 with the basic residues, bridges the TM helices thereby restricting the movement of the receptor in the intracellular regions. However, in the DDM environment, in the R and R’ states, although DDM head group forms hydrogen bonds with basic residues in the intracellular regions (Figs. 4J and 4K), they are not as strong as the salt bridges and do not bridge the helices and hence do not restrict the flexibility of the TM helices (Fig. 4C). The residues K7.56, K6.35, R5.66, R222 on ICL3 and R293 in the C-terminus of the receptor are involved in forming salt bridges with PIP2. This increased PIP2 interaction with the intracellular regions of the receptor is also borne out by the calculated non-bonded interaction energy between PIP2 and receptor in R and R’ states shown in Fig. S4.

### PIP2 staples the GPCR:G protein complex

PIP2 in the R*.G state acts as a “stapler” between the active state A_2A_R and the mini-Gs protein in the R*.G complex. The hydrophobic tail of the PIP2 interacts with the hydrophobic regions of the TM helices in R*.G state, and the charged head group of PIP2 makes salt bridge contacts with Arg/Lys residues in the miniGs protein (Fig. 4F). As shown in Fig. S4, this leads to a much stronger interaction energy of PIP2 with the R*.G complex compared to its interaction energies with R and R’ states of the receptor. Thus, PIP2 increases the stability of the R*.G complex by interacting with both these moieties. Such stapling interactions that strengthens the A_2A_R-miniGs complex were not observed in DDM (Fig. 4L).

### Intracellular calcium ions ease the receptor stiffening by PIP2 to modulate the basal activity of GPCRs

Cations as such as Ca^2+^ are known to play an important role as allosteric modulators of GPCR activity in cells (23–26). In our MD simulations in cell membrane with calcium, the Ca^2+^ ions make strong salt bridges with the phosphate groups of PIP2 resulting in breaking the PIP2 salt bridges with the intracellular Lys and Arg residues in the R and R’ states (Figs. 4G and 4H). Ca^2+^ coordinating with PIP2 thus frees up the Arg/Lys residues in the receptor. Consequently, the interaction energy between A_2A_R and PIP2 is weakened by Ca^2+^ ions and this trend is true for R, R’ and R*.G states (Fig. S4). This results in an increase in the flexibility of the TM helices which facilitates a few transitions between R and R’ states that was not observed in the cell membrane without Ca^2+^ (Fig. 4B). Based on this data we speculate that the localized concentration of PIP2, and Ca^2+^ in lipid rafts together modulate the basal activity of the GPCRs. Calcium ions also weaken the stapling interactions of PIP2 on the A_2A_R-miniGs complex (Fig. 4I). In an NMR study of the basal activity of β2-adrenergic receptor Staus et al speculated that the detergent as well as lipid nanodisc environments promote more transitions among the R, R’ and R*.G states compared to cell membrane(15). Thus, our findings provide a mechanism by which cell membranes restrain receptor conformations and thus modulate their basal activity. Calcium weakens the interaction energy of PIP2 with the receptor in all the three conformation states including the R*.G state (Fig. S4).

### Branched detergents mimic the behavior of PIP2 compared to unbranched detergents

In a previous study we examined the differences in the dynamic behavior of GPCRs in branched detergent micelles from the maltose-neopentylglycol (MNG) series compared to their unbranched counterparts such as DDM(30). We observed that LMNG tucks its branched hydrophobic tails between the TM helices in the intracellular region of the GPCR, while its two branched polar head groups make hydrogen bonds with the polar and basic residues in the intracellular region (Fig. S5A). The two head groups in LMNG being covalently linked through a central carbon atom aids the formation of the bifurcated hydrogen bonds and reduce the flexibility of the receptor in the intracellular region. This behavior of LMNG detergent is a mimic of the role of PIP2 in the cell membrane although the bifurcated hydrogen bonds are not as strong as the salt bridges made by PIP2 with the Arg/Lys in the intracellular part of the TM helices (Fig. S5B).

### Spatiotemporal heat map of GPCR activation microswitches reveal rheostat behavior rather than binary switches in cell membrane

A comprehensive analysis of the change in the inter-residue distances upon activation was performed by Zhou et al by comparing several inactive state and active state static structures of multiple class A GPCRs(10). This study elucidated the inter-residue distances (Fig. 5A and 5B) that show a distinct change going from inactive to active states and are known as activation “microswitches”. Comparison of static structures reveal a binary view (“on” or “off” states) of the activation microswitches. However, in the cell membrane environment the receptor exists in an ensemble of conformational states and the activation microswitches show differences in persistence frequency that calls for a continuum view of activation microswitches as in a *rheostat*. In order to examine this concept, we calculated the spatio-temporal heat map of the persistence frequency (percentage of MD snapshots showing these contacts made or broken within a cutoff distance – see Methods) of the activation microswitches made or broken from the MD simulation trajectories in cell membrane and in DDM detergent. As seen in Fig. 5B, there are four layers across the GPCR structure in which the activation microswitches are distributed. Layer 1 which is below the agonist binding site, involves the PIF motif, CWxP motif and the sodium binding site. The major conformational changes upon activation in this layer are the collapse of the sodium binding site (D^2.50^, S^3.39^ and N^7.45^) and repacking of P^5.50^ and F^6.44^ in the PIF motif (P^5.50^, F^6.44^ and I^3.40^). Layer 2, which is one step further down towards the intracellular region involves the opening of hydrophobic lock of L^3.43^, V^6.40^ and V^6.41^. Layer 3 involves the rewiring and compression of residues R^3.50^ and I^3.46^ and also expanding L^6.37^ and R^3.50^. Another import conformational change is the collapse of Y^7.53^ of the NPxxY motif, to its nearby residues L^3.43^, I^3.46^ and R^3.50^. Layer 4 which is close to the G-protein binding site undergoes releasing of R^3.50^ from D^3.49^ in the DRY motif and A^3.53^ motif upon activation. All the microswitches are in their active state in R*.G state in both detergent and in cell membrane with or without calcium (Fig. 5E). The interesting differences are in the heat map for NECA bound inactive state R and active-intermediate state R’. In the inactive R state, more activation microswitches in layer 1 close to the NECA binding site, and also microswitches centered at Y^7.53^ in layers 3 and 4 are active in detergent environment. Comparison between cell membrane with and without calcium, the presence of calcium improves activation around Y^7.53^ and around F^6.44^, similar to detergent micelle but to a lesser degree. In the cell membrane with and without calcium, the activation microswitches are active only in layer 1 close to NECA and not in layers 3 and 4 as in the detergent. The activation of microswitches centered at F^6.44^ in layer 1 and centered at L^3.43^ in layer 2 are active in the R state in detergent. All the three environments show similar activation profile in the R’ state with exception of PIF motif that shrinks more and I^6.40^ moves away from N^7.49^ in detergent micelle compared to cell membrane. Cell membrane with and without calcium show similar activation profile in R’ state. The shrinking of the sodium binding site typified by the distance D^2.50^ - S^3.39^, is not complete in cell membrane without calcium, while calcium enhances the extent of this shrinking and detergent micelle in the R state. Taken together, these results show that activation microswitches indeed function as rheostats. Cell membrane conditions limit the flexibility of the activation microswitches thus reducing the basal activity of the receptor. On the other hand, the detergent environment increases the flexibility of the receptor. While adding Ca^2+^ to cell membrane environment increase the receptor flexibility thus tuning some of the microswitches earlier than others, but to a less degree than detergent.

**Figure 5:**
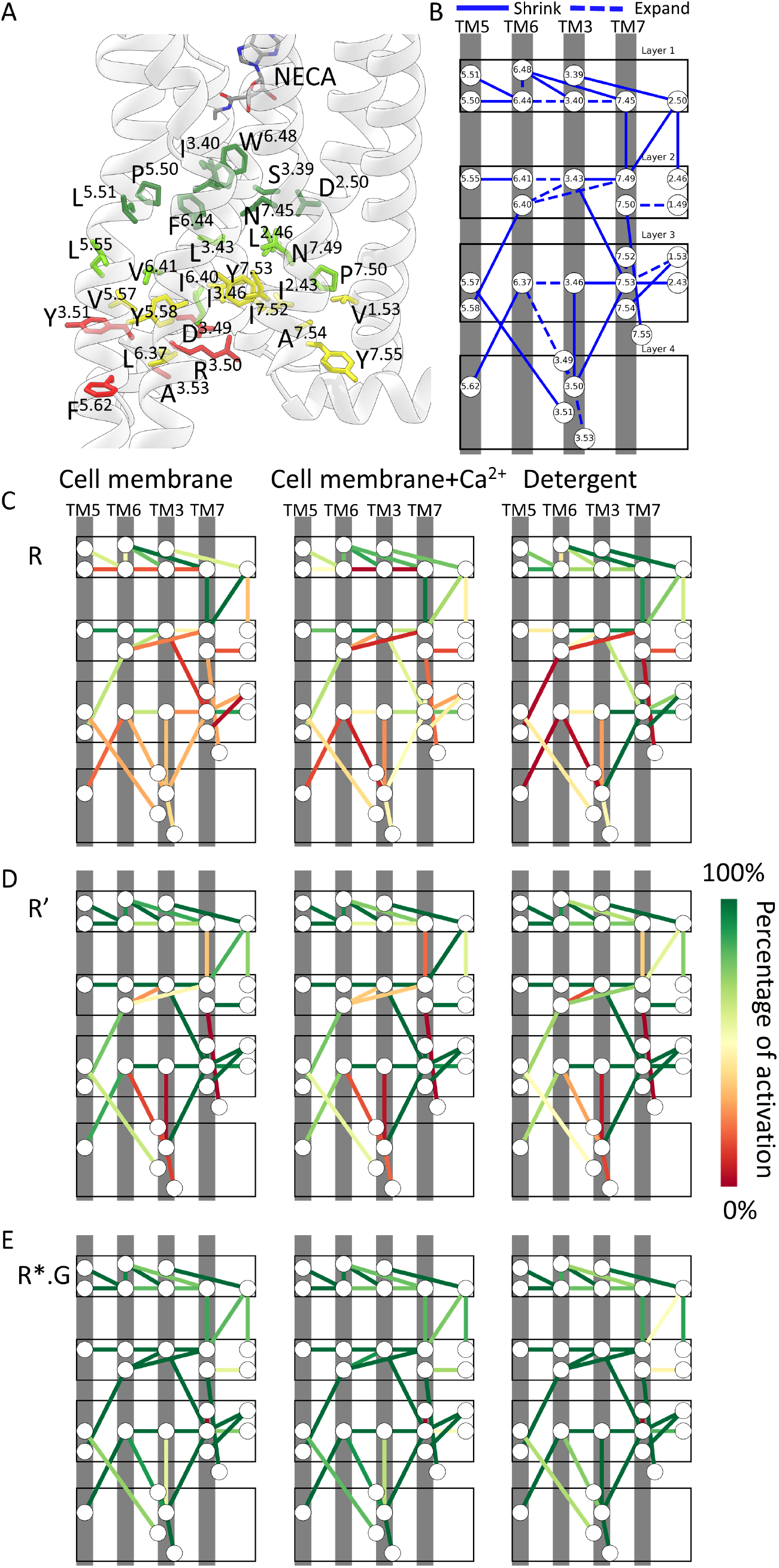
Activation microswitches behave like rheostats in cell membrane. **A.** Positions of residues labeled as activation microswitches by analysis of active and inactive state crystal structures. The agonist NECA is shown in grey, red and blue stick representation for reference. The microswitch residues are colored differently based on their positions relative to ligand binding site. Residues in layer 1 are colored in dark green (close to the agonist binding site), layer 2 in light green, layer 3 in yellow, layer 4 in red. **B.** A schematic showing the location of the inter-residue distances that are used as activation microswitches, and their respective TM helices shown as grey bars. The residue positions are shown by their Ballesteros-Weinstein numbering scheme commonly used for class A GPCR residue numbering. The solid line indicates that this inter-residue distance decreases upon activation, and the dashed line indicates the expansion of the inter-residue distance upon activation. **C.** The spatio-temporal heat map of the activation microswitches in the inactive R state in cell membrane, cell membrane with calcium and in DDM detergent. The heat scale from red to green shows the percentage of MD snapshots where each of the inter-residue distance is in the active state (see Methods for more details). **D-E**. The spatio-temporal heat map of the microswitches in the three environments for the R’ state (D) and R*.G state (E).

### Cholesterol binding sites

The persistence frequency of contacts between cholesterol and A_2A_R residues within a cutoff distance of 4.5Å were calculated from MD trajectories in cell membrane for all of R/R’/R*.G states. The persistence frequency for each residue is shown colored in a gray scale heat map on A_2A_R structure in Fig. S6. We identified three cholesterol binding sites: the first site is located in the extracellular region between TM1, TM2, the second site is located in the intracellular region around TM2, TM3 and TM4, and the third site is located in the extracellular region around TM6 and TM7. The first and third binding sites were also observed in the A_2A_R crystal structure 4EIY, where the binding site was lined with residues L^2.56^, F^2.60^, G^3.24^, C^3.25^, F^3.27^, I^3.28^, F^6.57^, I^6.53^, and L^6.49^ (9). The cholesterol binding site in the crystal structure was also observed in our simulations with >50% persistence contacts with all the residues listed above except G^3.24^, C^3.25^, F^3.27^ and I^3.28^. The second binding site located in TM2 and TM4 intracellular region was similar to the binding site identified in β2-adrenergic receptor crystal structure (pdb id:3D4S) (36). The residues lining this binding site are I^1.47^, L^1.48^, I^2.51^, W^4.50^, I^4.48^, I^4.45^ and V^4.51^. We observed cholesterol in this binding site with persistence frequency greater than 70% for the key residue contacts. The cholesterol binding sites are relatively conserved in all of R/R’/R*.G states, but they show differences in the persistent frequency of the contacts. The R’ state has highest persistence frequency with cholesterol in the TM1, TM2 site as well as the TM6, TM7 site. In the R*.G state, cholesterol binding site in the TM4, TM5 region is more persistent than the other two sites. In all the three states, TM3 has lowest persistence frequency with cholesterol, especially for R*.G state.

## Discussion

Although it is known that lipids and cations in the cell membrane play a functional role in modulating GPCR activity (16–18), the mechanism by which lipids and cations together modulate the receptor activity in comparison to detergents is unknown. We have used multi-scale MD simulations to decipher the mechanistic insights into the role of lipids and cations present in cell membrane on the flexibility and activity of agonist NECA bound A_2A_R in three different conformation states, R, R’ and R*.G. We have compared the dynamics of A_2A_R in these three states in cell membrane to that in DDM and LMNG detergent micelle environment that are commonly used for biophysical studies on GPCRs.

In this study, we observed that PIP2 accumulates close to the receptor in all the three R, R’ and R*.G states of A_2A_R but, play a different role in how they modulate the receptor activity and stability in each conformational state. PIP2 forms bifurcated salt bridges with Arg/Lys in the intracellular regions of the receptor in the R and R’ states. These bifurcated salt bridges act as interhelical clamps holding the TM helices in place thus making them less flexible and more stable in membrane. This reduces the number of transitions between inactive state and active-intermediate state in cell membrane as indicated by the yellow traffic light in Fig. 6. Presence of intracellular Ca^2+^ ions in the cell membrane breaks the salt bridges of PIP2 with the Arg/Lys in the TM helices and in the intracellular loops. This loosens the interhelical rigidity and promotes transitions between R and R’ states (green traffic light in Fig. 6). These observations are in line with the enhanced exchange rate between active and inactive conformations observed in NMR experiments(15, 22) in HDL nanodiscs and detergents. These authors further speculated that the lower level of basal activity in cells could come from cellular mechanisms that control basal activity. Here we provide insights into how the cell membrane components such as PIP2 and calcium maintain a balance between flexibility and stability in membranes. The bifurcated interactions that reduce the interhelical flexibility were also observed in branched detergents such as LMNG and not in unbranched detergent DDM. However, the stiffness provided by these bifurcated hydrogen bonds in LMNG are not as strong as the bifurcated salt bridges with PIP2. In the G protein bound fully active state of A_2A_R, PIP2 acts as a stapler between the A_2A_R and the miniGs protein thus enhancing the stability of the R*.G complex in its fully active state.

**Figure 6:**
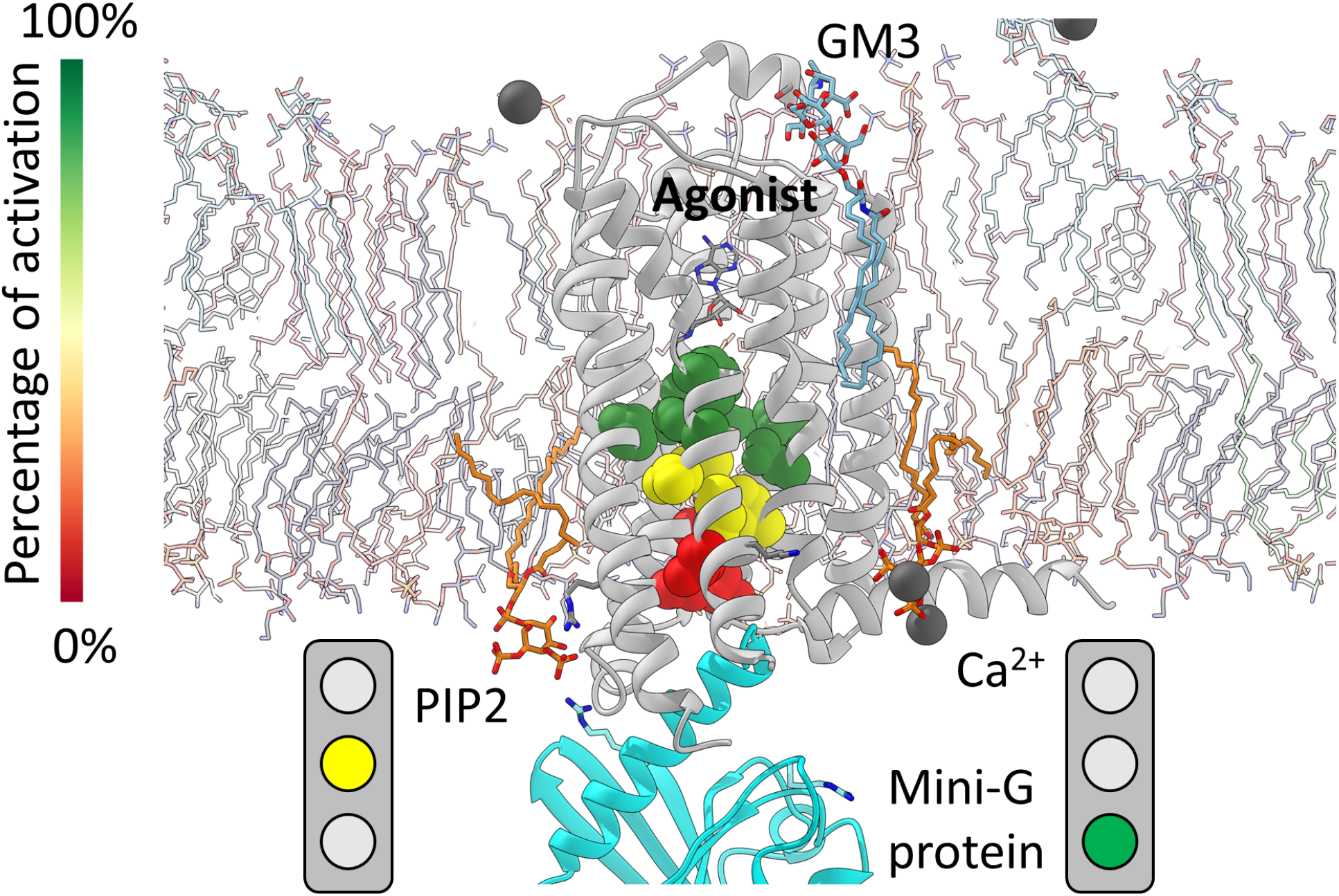
A model summarizing the findings from this study. GM3 shrinks the receptor-NECA binding site and enhances the binding affinity of agonist even without the G protein. PIP2 reduces the receptor flexibility and enhances stability in membranes showing a yellow light for receptor. Influence of calcium upsets the PIP2 interaction with the receptor thus increasing the receptor flexibility and activity shown by the green traffic light. The activation microswitches located at different layers from the agonist binding site show different levels of activation. This shows that activation functions like rheostats and not like binary switches.

Although experiments have shown that the binding affinity of the agonist NECA is an order of magnitude better in cells compared to detergents, there was no structural basis for this observation. Our ligand free energy calculations recapitulated this difference in binding affinity and showed that the enhanced binding affinity of NECA in cell membrane is due to the effect of GM3 enhancing the agonist receptor interactions and shrinking the NECA binding site, that does not happen in the detergent DDM environment. We also observed that GM3 pulls the extracellular loops down closer to the ligand binding site. Such strong modulatory interactions are absent in the detergent micelle. These differences between detergent and cell membrane environments on GPCR activity could yield different structural and biophysical properties when studied in detergents.

Our study shows that activation microswitches involved in GPCR activation function as rheostats rather than binary on and off switches as thought previously based on comparison of static structures. The spatio-temporal heat map of persistence frequency of the activation microswitches have shown that the lipid components of the cell membrane turn on different combination activation microswitches in the agonist bound intermediate R and R’ states. The propagation of the activation microswitches in different layers of the GPCR structure is different in cell membrane compared to detergents. MD simulations play a unique and critical role in delineating such mechanistic insights in modulating the differential effects of these activation microswitches compared to detergents.

In summary, our work exemplifies the role of GPCR structural dynamics and the effect of persistence frequency of lipids, cations or detergent interactions with the receptor in modulating the activity of GPCRs. The direct comparison between the cell membrane and detergent conditions allows us to draw insights into how cell membrane modulates the receptor activity in some cases similar to the detergent and many other cases differently from the detergent environment. These insights directly provide clues to design detergents or identify additives to detergent solutions that could better mimic the cell membrane properties. The observation that the components of the cell membrane constrain the receptor in various conformational states shows the synergy between sequence evolution and the endogenous environment that together provide the balance between the stability and the activity for GPCRs.

## Methods

### Coarse Grain (CG) simulation followed by All atom MD simulations

The workflow combining coarse grain MD with all-atom MD simulations described in this section is shown in Fig. S7. To study the effect of multiple lipids on the GPCR conformation ensemble, we used mixed lipid bilayer to mimic the cell membrane. The mix-bilayer membrane was generated using CHARMM-GUI,^52,53^ coarse grain (CG) membrane builder program.^54,55^ The outer leaflet contains POPC/DOPC/POPE/DOPE/sphingomyelin(Sph)/ganglioside(GM3)/cholesterol (Chol) lipids in ratio of 20:20:5:5:15:10:25, while the inner leaflet contained POPC/DOPC/POPE/DOPE/POPS/DOPS/phosphatidylinositol 4,5-bisphosphate(PIP2)/Chol in ratio of 5:5:20:20:8:7:10:25 and all simulations were neutralized by adding 0.15M of NaCl. The 50 nm × 50 nm simulation box has nine GPCR units that cover three of each agonist bound inactive state, active-intermediate state and G-protein bound fully active state. The nine units are placed 10~12 nm apart. The lipid bilayer was built three times to get statistically significant random distribution of lipids. After equilibration we performed 3*10μs of coarse grain MD (CGMD) simulations with Martini2.2 forcefield (37). We extracted 5 different lipid raft configurations from the coarse grain simulations for each conformational state of the receptor (R, R’ and R*.G). We chose those lipid configurations that remained a monomer (without dimerization) during the coarse grain MD simulations. The lipid environment surrounding the GPCR units were cut out into a 12 nm × 12 nm box. The CG models were converted to all-atom models using the script “backward.py” from the Martini website.(38) The original crystal structures (pdb code of 4EIY for R, 2YDV for R’ and 5G53 for R*.G) were placed in this equilibrated mixed lipid bilayer from coarse grain MD simulations, and solvated using a water box and neutralized with 0.15M of NaCl or 0.15M of CaCl_2_. The thermostabilizing mutations in crystal structure was converted back to the wild type residues using Maestro9.2, and disulfide bonds were built accordingly. The minimization-heating-equilibration-production was carried out using the protocol used in the previous work(39–42). Each of the 15 extracted all-atom simulation boxes were minimized and equilibrated using 50 ns long NPT equilibration simulations performed in steps to gradually reduce constraints on protein heavy atoms and lipid heavy atoms from 5 to 0 kcal/mol by 1 kcal/mol interval per 10 ns simulation windows. We performed 5 production runs with different starting velocities for each of R, R’ and R*.G states. Each production run, 400ns was run with NPT ensemble at 310 K with 2 fs time step using GROMACS with CHARMM36FF(43). Thus, we generated 2 μs of all-atom simulation for each system. This was done for each of the three conformational states in two environments namely: cell membrane, cell membrane with calcium. A total of 20 μs all-atom MD simulation have been done. We repeated this procedure for the antagonist ZMA241687 bound inactive state R (2 μs). We stored the MD snapshots every 20ps. For the all atom simulations the non-bond interactions were calculated with cutoff of 12 Å, particle mesh Ewald method was applied for van der Waals interactions calculation. The temperature was controlled by Nose-Hoover thermostat and pressure was controlled by Parrinello-Rahman method.

### CG simulation validation

Following the procedure outlined in reference(44), we validated the cell membrane coarse grain MD simulations using metrics such as: convergence of radial distribution function of different lipids, surface area per lipid in the top and bottom leaflet of the lipid bilayer to be within 1% (Fig. S8), also, the number of lipid molecules within 10 Å (first shell) of GPCR remains stable (Fig. S9). The radial distribution function (RDF) was used a metric to identify the lipids that accumulate close to the GPCR during the coarse grain simulations. RDF was calculated using vmd, as a function of the distance between mass centers of GPCR TM region and heavy atoms of each lipid. The counting of the number of lipids counting was done for every type of lipid within 10 Å of the mass center of GPCR TM region. Area per lipid is calculated using membrane plugin, a plugin in VMD by choosing the head group beads from CG model.(45) The coarse grain MD simulations showed clustering of cholesterol, PIP2 and GM3 close to the receptor in all the three conformation states (Fig. S1A), as evident from the radial distribution function and the number of each type of lipid calculated around the receptor (see Fig. S1B and S1C of the Supporting Information).

### All atom MD simulations in detergent DDM environment

We have performed the MD simulations of wild type of inactive state (R), active-intermediate (R’), and fully active state of A_2_AR under DDM detergents using GROMACS package with CHARMM36FF(43, 46). The initial coordinates of GPCR with the agonist NECA were taken from the PDB code of 2YDV and 5G53, respectively. All thermostable mutants (L48A^TM2^, A54L^TM2^, T65A^TM2^, Q89A^TM3^, and N154A^ECL2^) were mutated back to the wild type sequence using Maestro9.2. We used 192 DDM molecules to build each receptor-micelle complex using CHARMM-GUI/Micelle Builder module. We used the TIP3 force field for describing the waters. The receptor-micelle complexes were equilibrated using the same MD simulations as described in above mixed-bilayer membrane systems. We performed 5 production runs with different starting velocities each 400ns long. The trajectories of the all-atom MD simulations in the LMNG series of detergents were taken from our previous work(30) and used here for comparison. We used the TIP3 force field for describing the waters. All the analysis described below we took the second half of each trajectory for each state (R, R’, and R*.G) and each environment (cell membrane, cell membrane with calcium, detergent) from 200ns to 400ns and concatenated them for 5 runs (totaling to 1 μs) to perform analyses described below.

### Inter-residue distance analysis

The distances between Cα atoms of residues 3.50 and 6.34, 3.50 and 7.53 shown in Fig. 4A-C, were calculated combining using the aggregated trajectories for each system. The distances are plotted as 2D heatmap using 80 bins after normalizing the population in each bin.

### Extracellular loop interactions with GM3 or DDM

The distance between all heavy atoms from residues located in the extracellular (ECL)loops and heavy atoms of GM3, for cell membrane system, or DDM system were measured. The interaction is considered made if any of the distance is smaller than 4.5 Å. The persistence frequency is the percentage of MD snapshots that contain a prescribed contact.

### Ligand binding pocket volume analysis

The trajectory for R state in both DDM and cell membrane were clustered by ligand RMSD after aligning the MD snapshots by GPCR TM region. The most populated top conformational cluster from both systems (cell membrane and DDM) was chosen and representative structure was used for binding pocket volume analysis. The software “*trj_cavity*” was used to calculate binding pocket volume, with 1.7 Å as probe radius (47). To calculate the fluctuation in the NECA binding site volume, we averaged the volume over 10 frames taken from trajectory at equal interval every 100 ns. In this analysis the average and standard deviation of ligand binding pocket volume in DDM is 237.2 ±36.4 Å^3^ and 383.6 ± 150.7 Å^3^ in cell membrane.

### Ligand binding pose and pocket residues flexibility analysis

The flexibility of the ligand was calculated using conformational clustering. We clustered the NECA conformations by aligning the MD snapshots by the receptor to the reference structure and then clustered based on the RMSD in coordinates of NECA heavy atoms. The trajectory of R state in both cell membrane and detergent micelle were taken for ligand binding pose clustering analysis. The trajectory was aligned base on TM region backbone atoms, then the ligand heavy atoms were used for the RMSD in co-ordinates calculation. DBscan method in MDanalysis(48) was used to cluster the ligand poses based on RMSD matrix with 1.5 Å as cutoff for each cluster. The GPCR structure backbone heavy atoms from trajectory were used for RMSF calculation for the receptor residues.

### Activation Micro-switch analysis

We extracted all the inter-residue pair distances that were characterized to undergo changes (either increase or decrease) upon activation(10). The distance between centers of mass of each residue pair was calculated. The cutoff distance for defining the inactive state of the activation microswitch distance was extracted from the MD simulations of antagonist bound inactive state R of A_2A_R. We calculated the distance distribution of all the pairs of activation microswitches. The distances histograms were fitted to a Gaussian function to extract the mean distance from the distribution and the standard deviation. We did not use a single distance extracted from a static structure, because under cell membrane conditions there is always a range of distances that is defined as inactive state. The mean value minus one standard deviation for the activation microswitch pairs that are known to shrink, was used as cutoff distance to separate active and inactive states. For those activation microswitch pairs that are known to expand upon activation, the mean plus one standard deviation was used as cutoff distance to define the active state. The same cutoff distance was applied to simulations in cell membrane and detergent micelle bound with NECA, for R, R’ and R*.G states. Once the cutoff was determined, the distance histograms for all microswitch pairs from all the simulations were generated. The population below the cutoff for the shrinking pairs (above the cutoff for the expanding pairs) were counted and the percentages over whole population were calculated to arrive at the percentage of activation.

### Interaction energy of PIP2 with the receptor

The interaction energy between PIP2 and the receptor was calculated as average of the sum of electrostatic energy and van der Waal interaction energy between all PIP2 molecules and all the residues in A_2A_R, averaged over the concatenated trajectory for each system.

### Cholesterol persistence frequency

We identified the cholesterol molecules within 4.5 Å of every A_2A_R residue and calculated the persistence frequency of these contacts as the percentage of MD snapshots that showed such contacts. The contact frequencies were set as B-factor for each residue and colored as heatmap on GPCR structure. This was also plotted as a bar graph for each of R/R’/R*.G states.

### Free energy perturbation method for calculating the Ligand binding free energy

The binding free energy (ΔG) of NECA in three conformational states (R, R’, and R*.G) and two environments (DDM micelle and cell membrane) shown in Fig. 2A was calculated by BAR (Bennett acceptance ratio) algorithm in the GROMACS package (49). BAR calculation was conducted at different values of λ (coupling parameter) from ligand-bound (λ=0) to ligand–free (λ=1) using equidistance spacing of 0.05. We performed 10ns of all-atom MD simulations for each λ value.

## Acknowledgements

Funding for this work was provided by NIH R01 grants GM097261 and GM117923 to N.V. We acknowledge insightful discussions with Dr. Supriyo Bhattacharya.

## Supporting information

**Figure S1.**
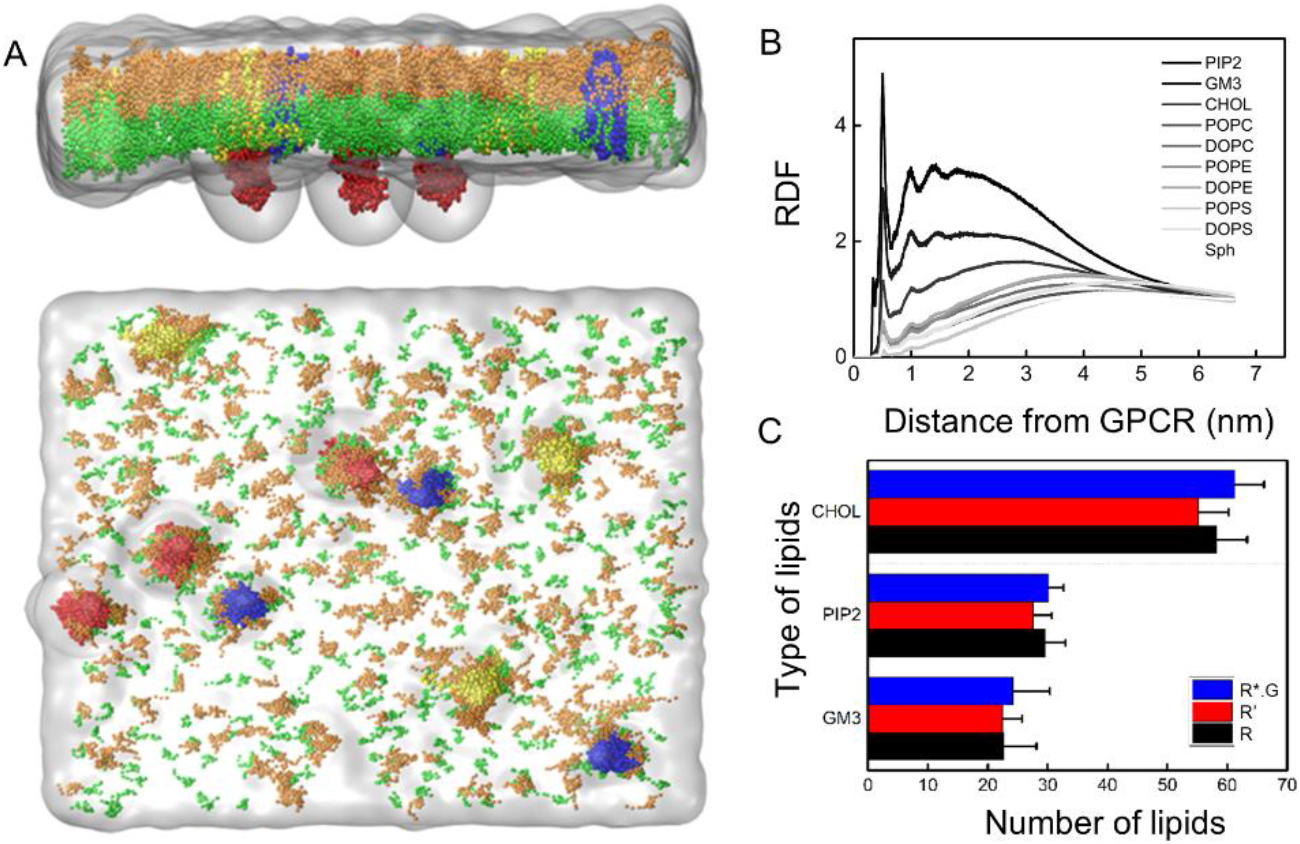
**A)** The clustering of GM3 (orange) and PIP2 (green) around the receptors (inactive R state of the adenosine receptor A_2A_R is shown in blue, active-intermediate state R’ in yellow, fully active R*.G state in red) in CG simulation. **B)** The radial distribution function RDF shows PIP2, GM3 and CHOL are the three lipids that cluster close to all the three conformational states (R, R’ and R*.G) of A_2A_R. **C)** Lipid count of PIP2, GM3 and Cholesterol near A_2A_R. The number of lipids in R/R’/R*.G states did not show significant difference.

**Figure S2:**
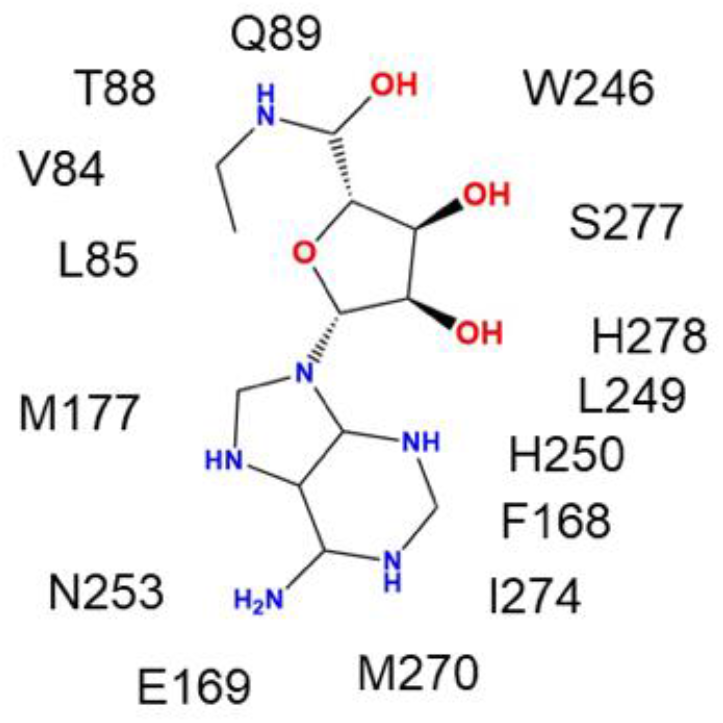
The residue contacts observed in the crystal structure of NECA bound to A_2A_R (pdb ID:2YDV). The persistence frequency for these contacts are shown in Figure 2B of the main text.

**Figure S3.**
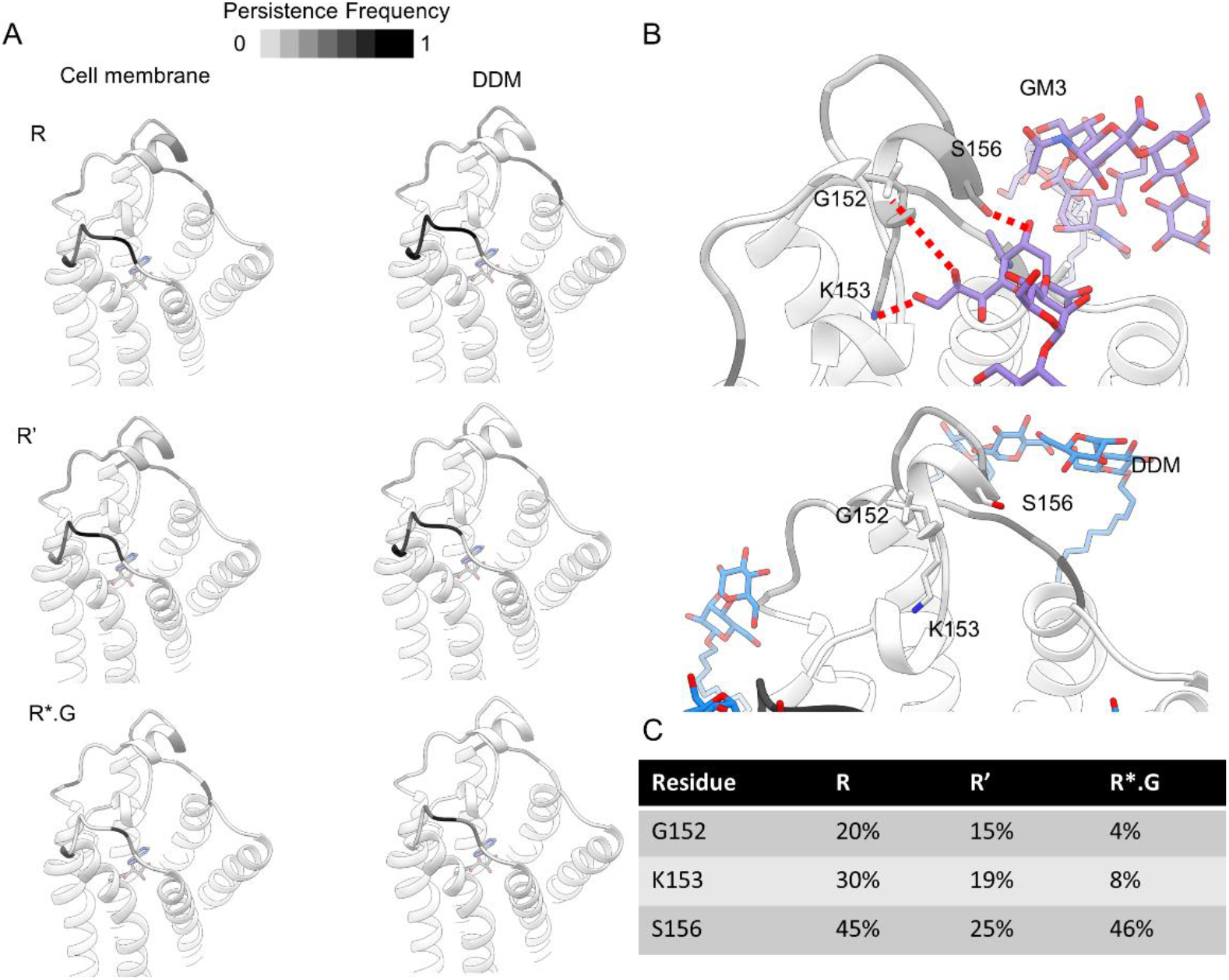
**A)** The residues in the extracellular loops ECL1 and ECL2 are shaded by the persistence frequency (percentage of MD snapshots) of interactions of GM3 head groups and with the heavy atoms in DDM. **B)** GM3 head group interactions with the three residues G152, K153 and S156 on ECL2 in both cell membrane and detergent. **C)** The persistence frequency of contact between important ECL2 residues and GM3 in cell membrane simulations for all the three states.

**Figure S4.**
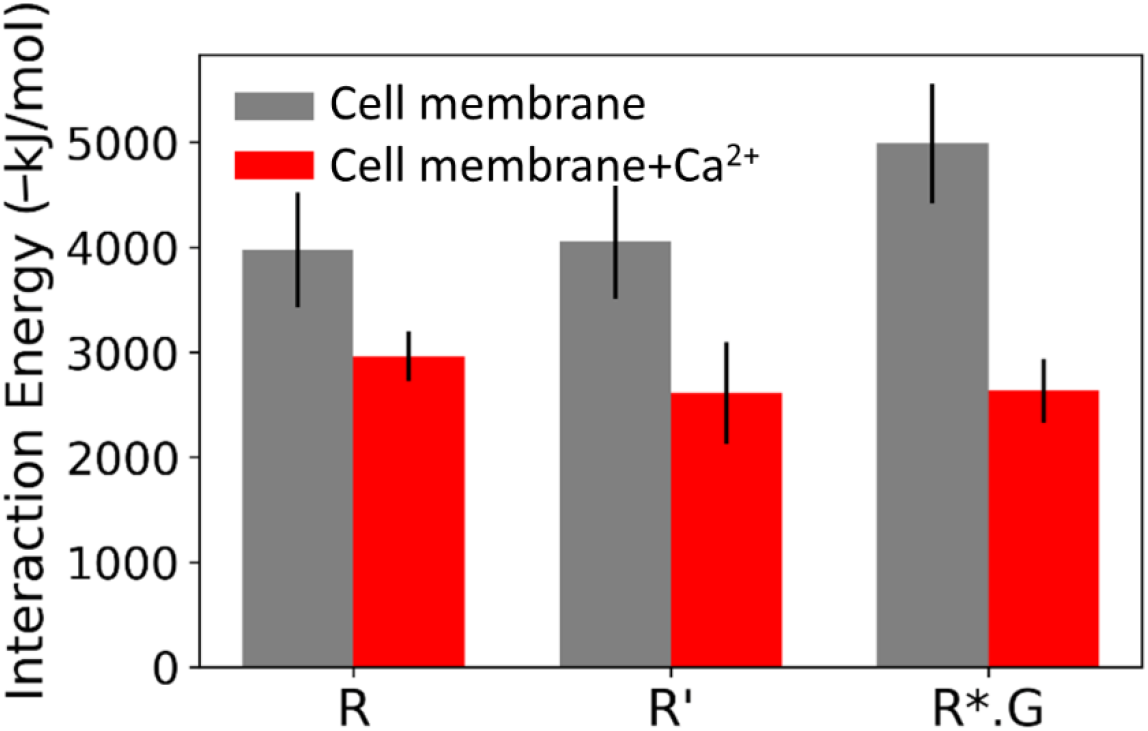
Non-bond interaction energy between PIP2 and all the residues in the receptor A_2A_R in various states in cell membrane with and without calcium. The energies are calculated as the average over the all-atom MD simulation trajectories (see Methods).

**Figure S5.**
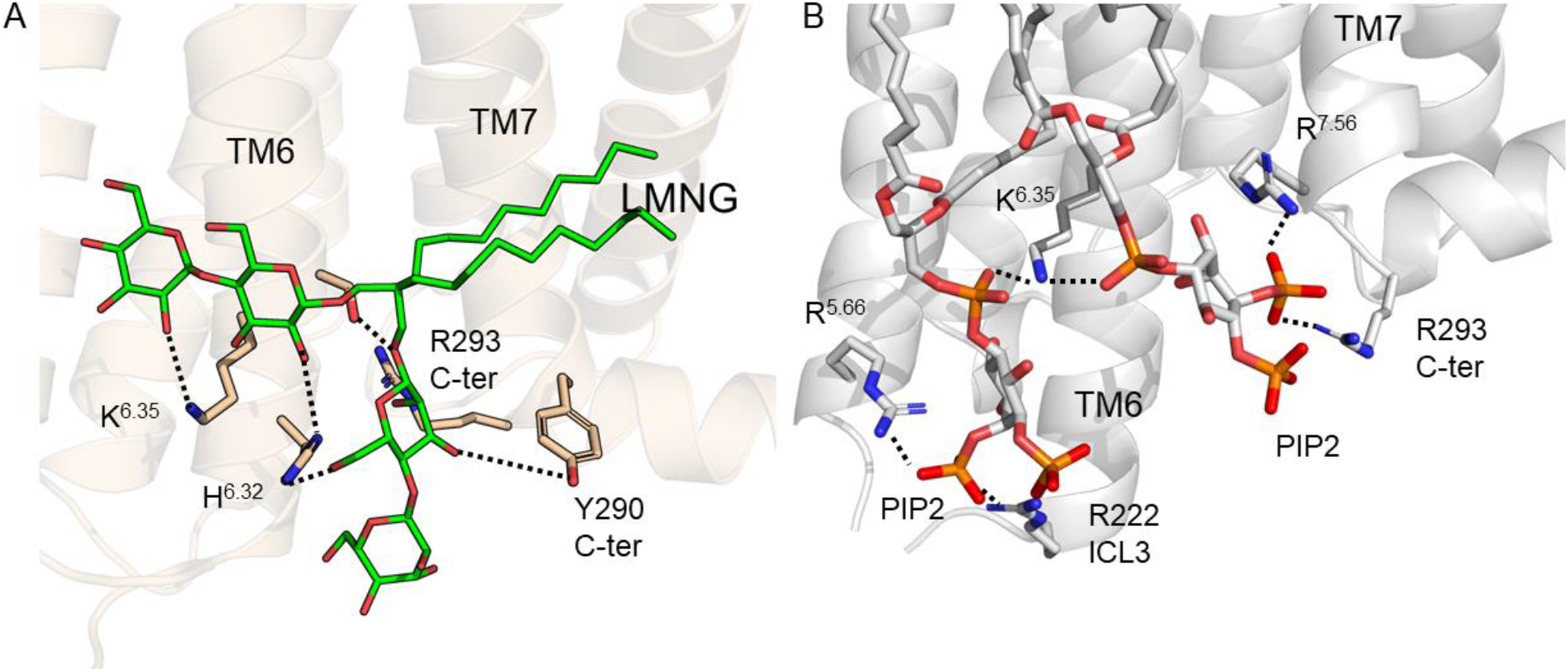
Branched detergent LMNG mimics PIP2 of the cell membrane. **A.** LMNG (shown in green and red sticks) interactions with the basic residues in the intracellular regions of TM6 and TM7 in A_2A_R in the R state. **B**. PIP2 bridging interactions with Arg/Lys in TM intracellular region of A2AR in the R state.

**Figure S6.**
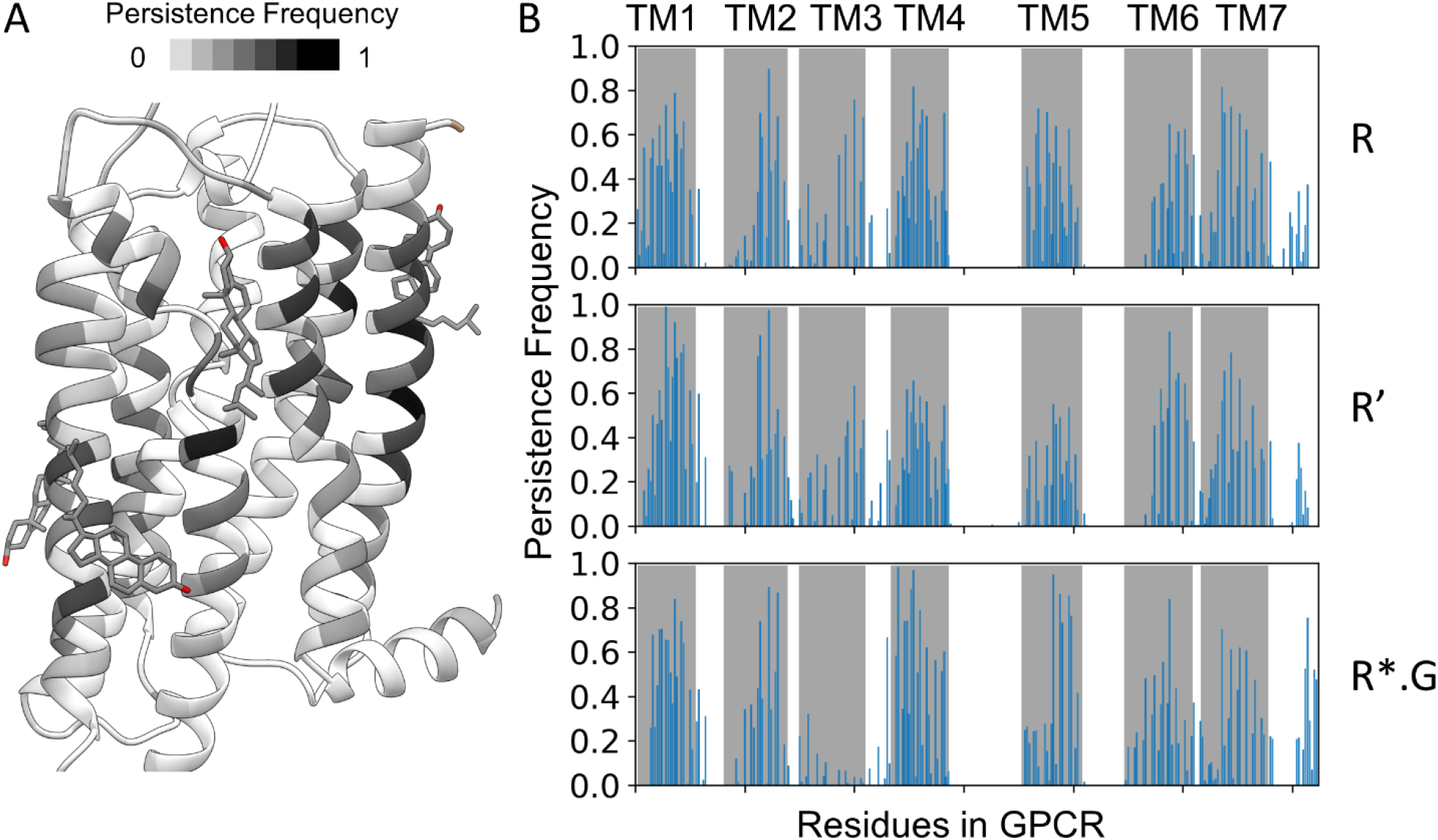
Cholesterol binding site identified by contact frequency shown as heat map (left) and frequency (right).

**Figure S7.**
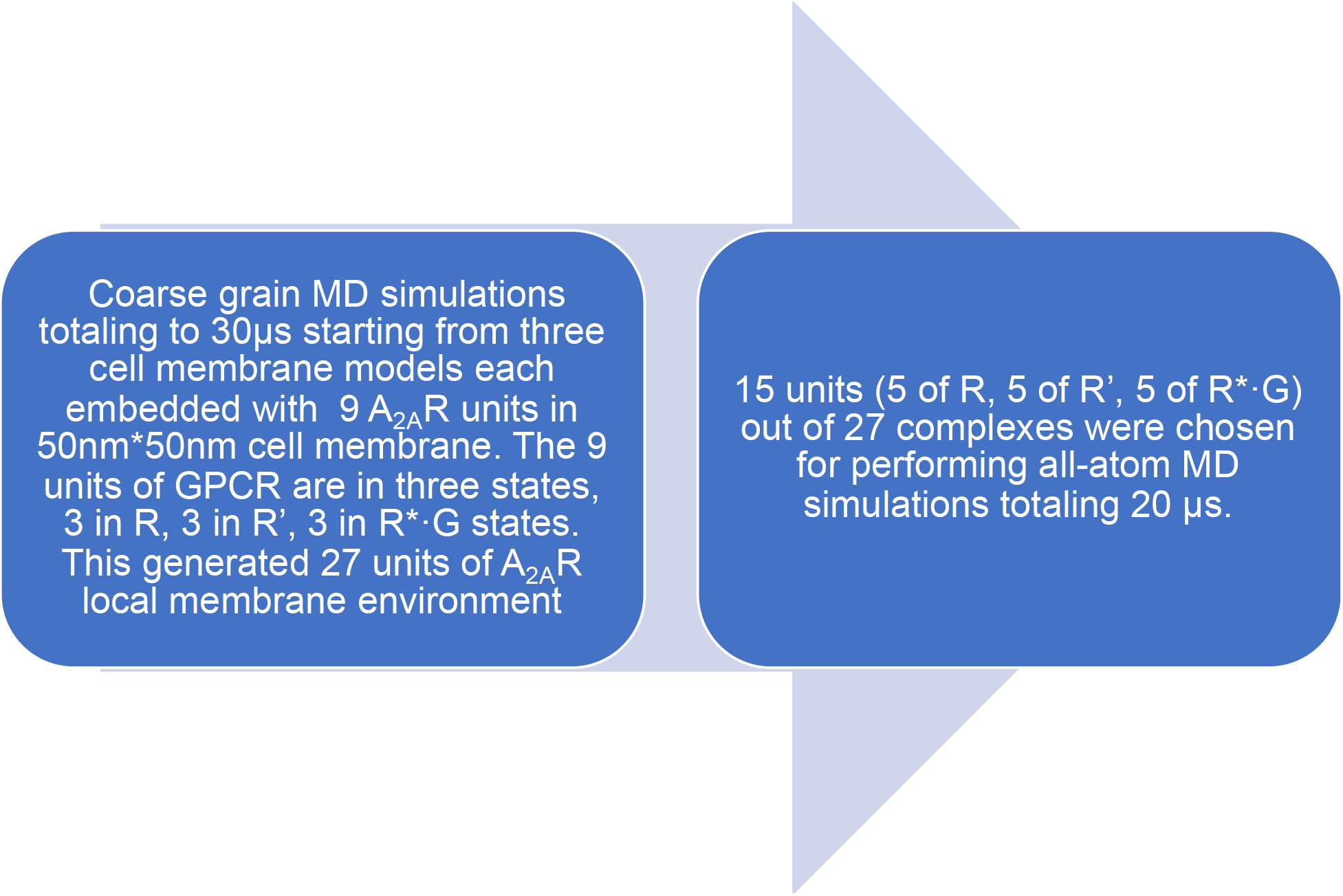
Flow chart of multi-scale MD simulation protocol used in this study.

**Figure S8.**
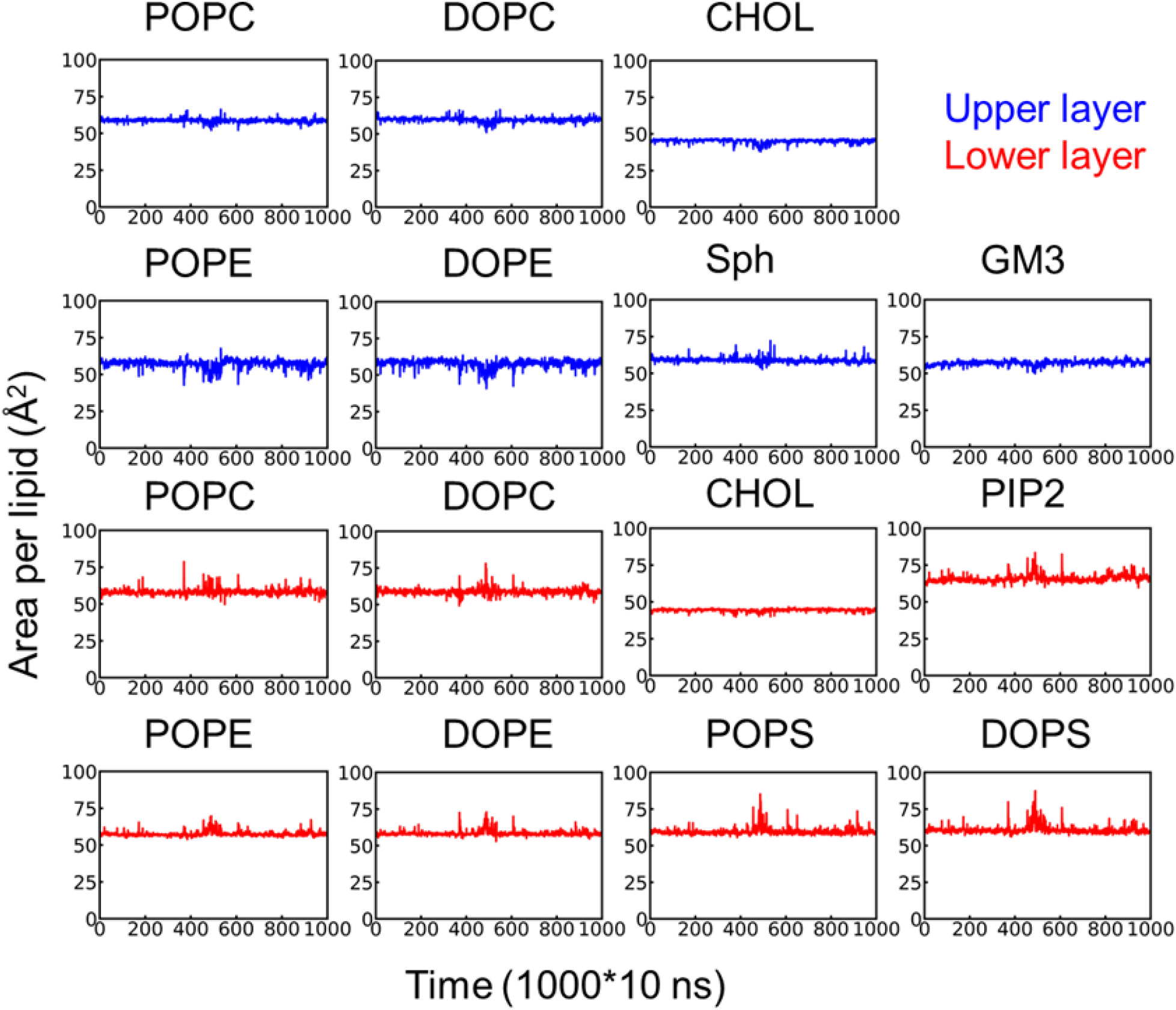
Area per lipid (APL) over CG simulation time (10 μs). The lipids in the outer leaflet of the membrane bilayer are shown in blue and inner leaflet in red. Each time window is 10 μs long.

**Figure S9.**
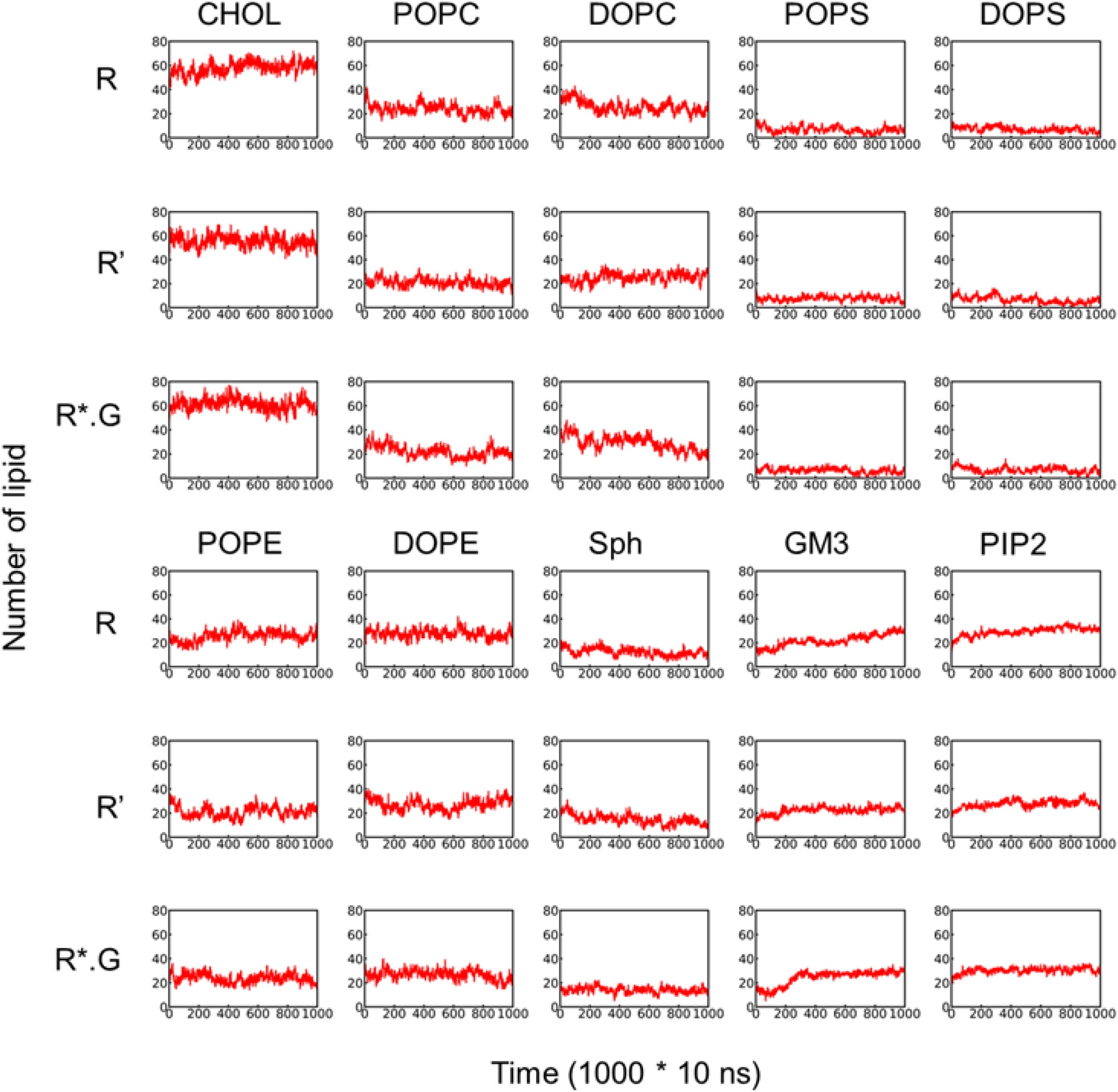
Number of each type of lipid molecules in first shell (within 10 Å of center of mass of the receptor of the TM region) surrounding GPCR over CG simulation time as a validation of equilibrium reached.

**Table S1.** Type of Lipids and their composition used in MD simulation to mimic cell membrane.

| Lipid | POPC | DOPC | POPE | DOPE | POPS | DOPS | Sph | GM3 | PIP2 | CHOL |
| --- | --- | --- | --- | --- | --- | --- | --- | --- | --- | --- |
| Upper layer | 20% | 20% | 5% | 5% | 0% | 0% | 15% | 10% | 0% | 25% |
| Lower layer | 5% | 5% | 20% | 20% | 8% | 7% | 0% | 0% | 10% | 25% |

**Table S2.**
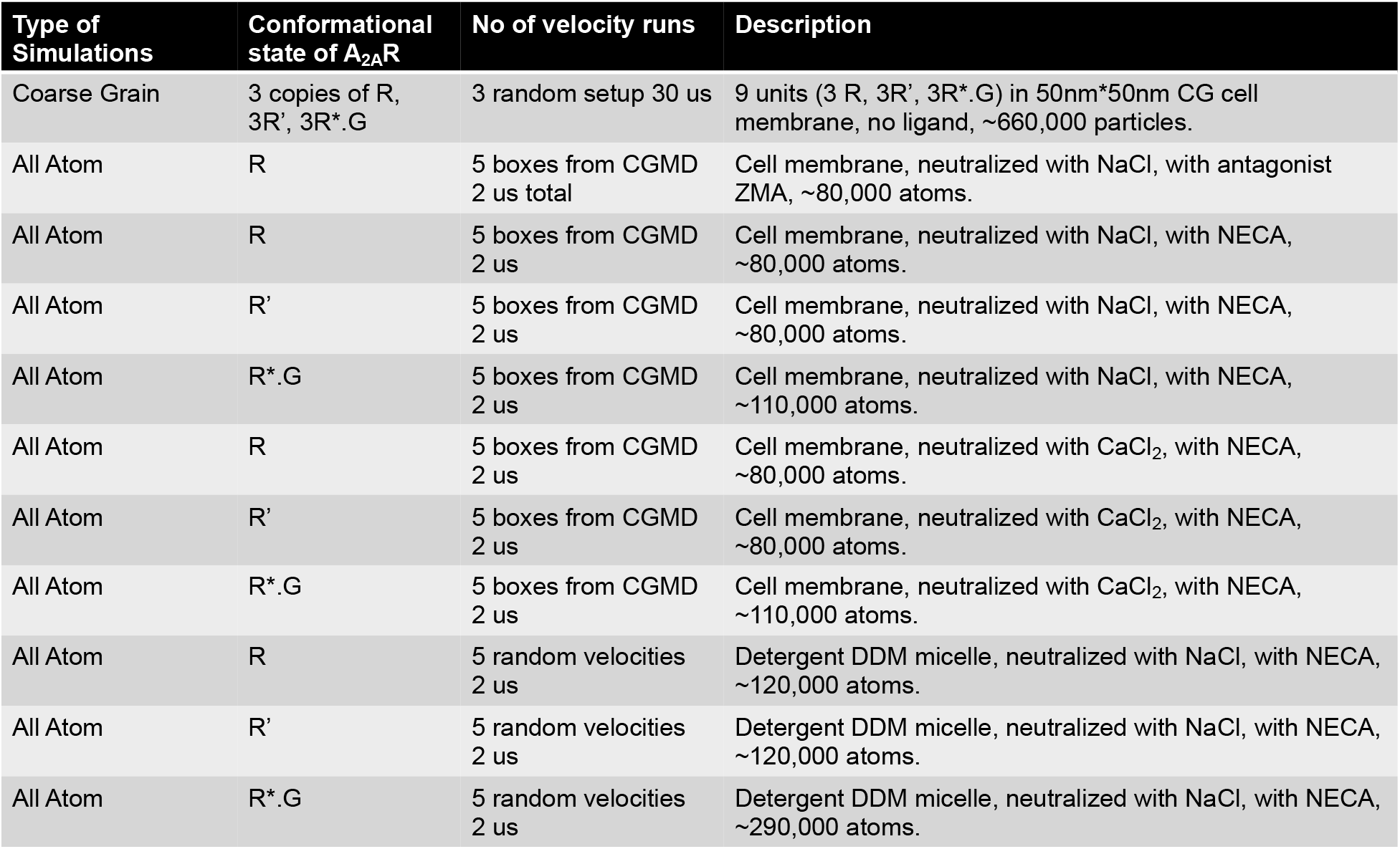
Details of all the simulations carried out in this study.

